# Evolution trajectories of snake genes and genomes revealed by comparative analyses of five-pacer viper

**DOI:** 10.1101/067538

**Authors:** Wei Yin, Zong-ji Wang, Qi-ye Li, Jin-ming Lian, Yang Zhou, Bing-zheng Lu, Li-jun Jin, Peng-xin Qiu, Pei Zhang, Wen-bo Zhu, Bo Wen, Yi-jun Huang, Zhi-long Lin, Bi-tao Qiu, Xing-wen Su, Huan-ming Yang, Guo-jie Zhang, Guang-mei Yan, Qi Zhou

## Abstract

Snake’s numerous fascinating features distinctive from other tetrapods necessitate a rich history of genome evolution that is still obscure. To address this, we report the first high-quality genome of a viper, *Deinagkistrodon acutus* and comparative analyses using other species from major snake and lizard lineages. We map the evolution trajectories of transposable elements (TEs), developmental genes and sex chromosomes onto the snake phylogeny. TEs exhibit dynamic lineage-specific expansion. And in the viper many TEs may have been rewired into the regulatory network of brain genes, as shown by their associated expression with nearby genes in the brain but not in other tissues. We detect signatures of adaptive evolution in olfactory, venom and thermal-sensing genes, and also functional degeneration of genes associated with vision and hearing. Many *Hox* and *Tbx* limb-patterning genes show evidence of relaxed selective constraints, and such genes’ phylogenetic distribution supports fossil evidence for a successive loss of forelimbs then hindlimbs during the snake evolution. Finally, we infer that the Z and W sex chromosomes had undergone at least three recombination suppression events at the ancestor of advanced snakes, with the W chromosomes showing a gradient of degeneration from basal to advanced snakes. These results, together with all the genes identified as undergoing adaptive or degenerative evolution episodes at respective snake lineages forge a framework for our deep understanding into snakes’ molecular evolution history.

---

Snakes have undergone a massive adaptive radiation with ~3400 extant species successfully inhabiting almost all continents except for the polar regions^1^. This process has culminated in ‘advanced snakes’ (Caenophidia, ~3000 species), involved numerous evolutionary changes in body form, chemo-/thermo-perception, venom and sexual reproductive systems, which together distinguish snakes from the majority of other Squamates (lizards and worm lizards). Some of these dramatic changes can be tracked from fossils, which established that the ancestor of snakes had already evolved a elongated body plan, probably as an adaptation to a burrowing and crawling lifestyle, but had lost only the forelimbs^2-4^. Extant boa and python species retain rudimentary hindlimbs, while advanced snakes have completely lost them. The limblessness, accompanied by degeneration in visual and auditory perception have not compromised snakes’ dominant role as top predators, largely due to the evolution of infrared sensing and/or venom, and the development of corresponding facial pit and fangs (specialized teeth for venom injection) independently in different lineages^5,6^.

These extreme adaptations have sparked strong and standing interest into their genetic basis. Snakes are used as a model for studying various basic questions like mechanisms of axial patterning and limb development^3,7,8^, ‘birth-and-death’ of venom proteins^9-11^, and also sex chromosome evolution^12^. Cytogenetic findings in snakes first drove Ohno to propose that sex chromosomes in vertebrates evolved from ancestral autosomes^13^, like those of insects^14^ and plants^15^. Our insights into these questions have been recently advanced by the application of next-generation sequencing. Analyses of python and king cobra genomes and transcriptomes have uncovered the metabolic gene repertoire involved in feeding, and inferred massive expansion and adaptive evolution of toxin families in elapids (an ‘advanced’ group)^10,16^. However, comparative studies of multiple snake genomes unraveling their evolutionary trajectories since the divergence from lizards are lacking, and so far only a few specific developmental ‘toolbox’ (e.g., *Hox*^7,17,18^ and Fgf signaling pathway^19^) genes have been studied between snakes and lizards. This hampers our comprehensive understanding into the molecular basis of stepwise or independent acquisition of snake specific traits. We bridge this gap here by deep-sequencing the genomes and transcriptomes of five-pacer viper (*Deinagkistrodon acutus*, Figure 1A), a member of the Viperidae family. This pit viper is a paragon of infrared sensing, heteromorphic ZW sex chromosomes, and distinctive types of fangs and toxins (its common name exaggerates that victims can walk no more than five paces) from other venomous snake families^6,20^. Despite the intraspecific differences within the same family, comparative analyses to the available genomes of three other major snake family species, i.e., *Boa constrictor* (Boidae family)^21^, *Python bivittatus* (Pythonidae family)^16^, *Ophiophagus hannah* (Elapidae family)^10^ and several reptile outgroups should recapitulate the major genomic adaptive or degenerative changes that occurred in the ancestor and along individual branches of snake families, and would also promote current antivenom therapy and drug discovery (e.g., thrombolytics) out of the viper toxins^22^.

**Figure 1.**
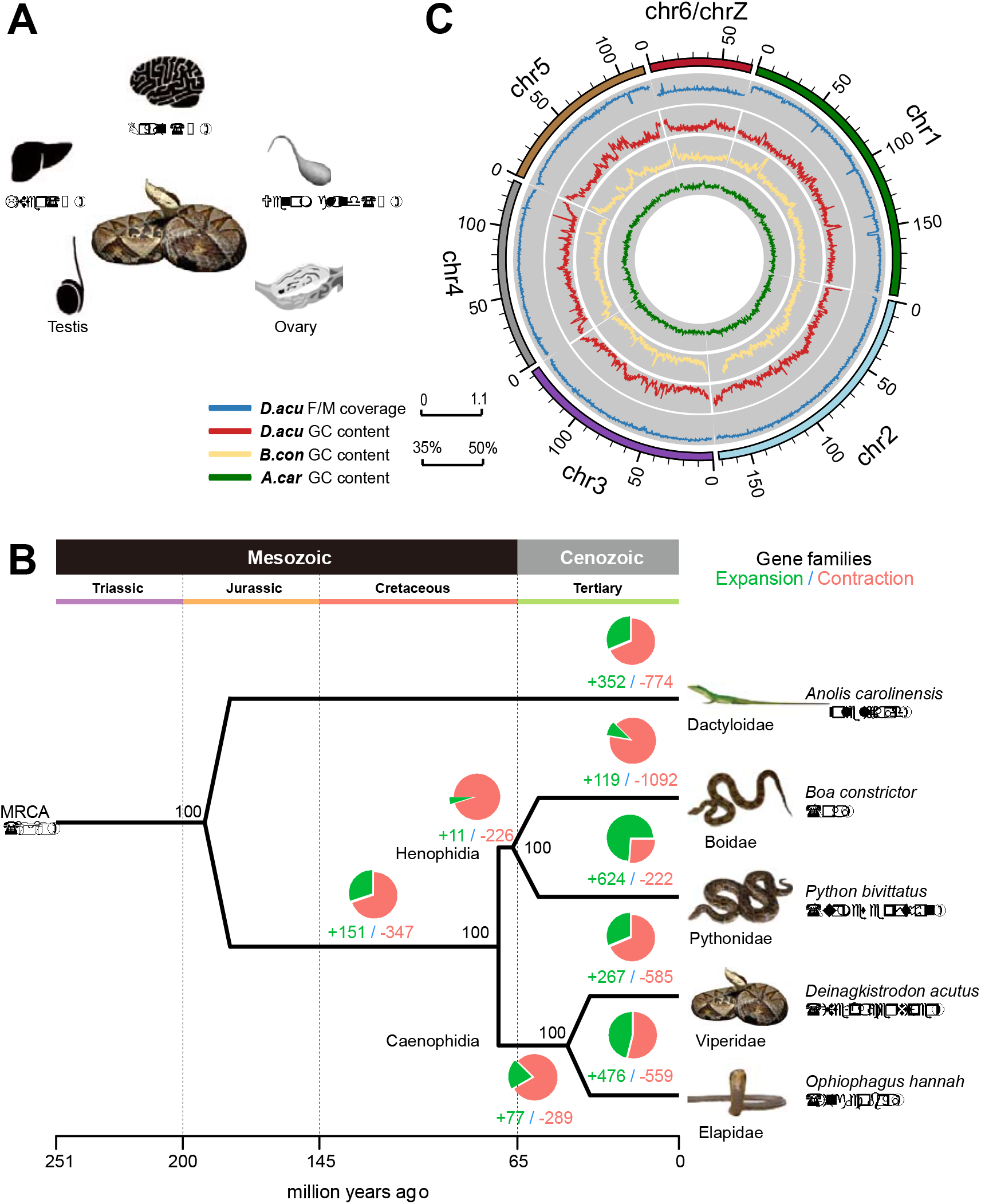
The comparative genomic landscape of five-pacer viper. **(A)** *Deinagkistrodon acutus* (five-pacer viper) and eight adult tissues used in this study. **(B)** Circos plot showing the linkage group assignment using lizard chromosomes as reference (outmost circle), normalized female vs. male mapped read coverage ratio (blue line) and GC-isochore structures of five-pacer viper (red), boa (yellow) and green anole lizard (green). Both snake genomes have a much higher variation of local GC content than that of green anole lizard. **(C)** Phylogenomic tree constructed using fourfold degenerate sites from 8006 single-copy orthologous genes. We also showed bootstrapping percentages, the numbers of inferred gene family expansion (in green) and contraction (red), and corresponding phylogenetic terms at each node. MRCA: most recent common ancestor.

## Evolution of snake genome architecture

We sequenced a male and a female five-pacer viper to extremely high-coverage (♀ 238 fold, ♂ 114 fold, **Supplementary Table 1**), and estimated the genome size to be 1.43Gb based on k-mer frequency distribution^23^ (**Supplementary Table 2, Supplementary Figure 1**). Fewer than 10% of the reads, which have a low quality or are probably derived from repetitive regions, were excluded from the genome assembly (**Supplementary Table 3**). We generated a draft genome using only male reads for constructing the contigs, and female long-insert (2kb~40kb) library reads for joining the contigs into scaffolds. It has an assembled size of 1.47Gb, with a slightly better quality than the genome using purely female reads. It has high continuity (contig N50: 22.42kb, scaffold N50: 2.12Mb) and integrity (gap content 5.6%, **Supplementary Table 4**), thus was chosen as the reference genome for further analyses. It includes a total of 21,194 predicted protein-coding genes, as estimated using known vertebrate protein sequences and massive transcriptome data generated in this study from eight tissues (Figure 1A, **Materials and Methods**). For comparative analyses, we also annotated 17,392 protein-coding genes in the boa genome (the SGA version from^21^). 80.84% (17,134) of the viper genes show robust expression (normalized expression level RPKM>1) in at least one tissue, comparable to the value of 70.77% in king cobra (**Supplementary Table 5**). Based on 5,353 one-to-one orthologous gene groups of four snake species (five-pacer viper, boa^21^, python^16^, king cobra^10^), green anole lizard^24^ and several other sequenced vertebrate genomes (**Materials and Methods**), we constructed a phylogenomic tree with high bootstrapping values at all nodes (Figure 1B). We estimated that advanced snakes diverged from boa and python about 66.9 (47.2~84.4) million years ago (MYA), and five-pacer viper and king cobra diverged 44.9 (27.5~65.0) MYA assuming a molecular clock. These results are consistent with the oldest snake and viper fossils from 140.8 and 84.7 MYA, respectively^25^.

The local GC content of snakes (boa and five-pacer viper) shows variation (GC isochores) similar to the genomes of turtles and crocodiles, and intermediate between mammals/birds and lizard (Figure 1C, **Supplementary Figure 2**), confirming the loss of such a genomic feature in lizard among tetrapods^24^. Cytogenetic studies showed like most other snakes, the five-pacer viper karyotype has 2n=36 (16 macro- and 20 micro-chromosomes) chromosomes^26^, with extensive inter-chromosomal conservation with the lizard^27^. This enables us to organize 56.50% of the viper scaffold sequences into linkage groups, based on their homology with sequences of known green anole lizard macro-chromosomes (**Supplementary Table 6**). As expected, autosomal sequences have the same read coverage in both sexes, whereas scaffolds inferred to be located on the viper Z chromosome (homologous to green anole lizard chr6) have coverage in the female that is half that in the male (Figure 1C). Additionally, the frequency of heterozygous variants on the Z chromosome is much lower in the female than in the male (0.005% vs 0.08%, *P*-value<2.2e-16, Wilcoxon signed rank test) due to the nearly hemizygous state of Z chromosome in female, while those of autosomes (~0.1%) are very similar between sexes. These results indicated that our assembly mostly assigns genes to the correct chromosome, and this is further supported by comparisons of 172 genes’ locations with previous fluorescence *in situ* hybridization results (**Supplementary Data 1**)^27^. They also suggest that the viper sex chromosomes are highly differentiated from each other (see below).

47.47% of the viper genome consists of transposable element sequences (TEs), higher than any other snakes so far analyzed (33.95~39.59%), which cannot be explained solely by the higher assembly quality than that of the other species genome sequences^10,16,21^ (**Supplementary Table 7-8**). These are mostly long interspersed elements (LINE, 13.84% of the genome) and DNA transposons (7.96%, **Supplementary Table 7**). Sequence divergence of individual families from inferred consensus sequences uncovered recent rampant activities in the viper lineage of LINEs (CR1), DNA transposons (hAT and TcMar) and retrotransposons (Gypsy and DIRS). In particular, there is an excess of low-divergence (<10% divergence level) CR1 and hAT elements in the viper genome only (Figure 2A). We also inferred earlier propagation of TEs shared by viper and king cobra, which thus probably occurred in the ancestor of advanced snakes. Together, these derived insertions resulted in an at least three-fold difference in the CR1 and hAT content between viper and more basal-branching snakes like boa and python (Figure 2B). While boa and python have undergone independent expansion of L2 and CR1 repeats, so that their overall LINE content is at a similar level to that of the viper and cobra (Figure 2A, **Supplementary Table 7**).

**Figure 2.**
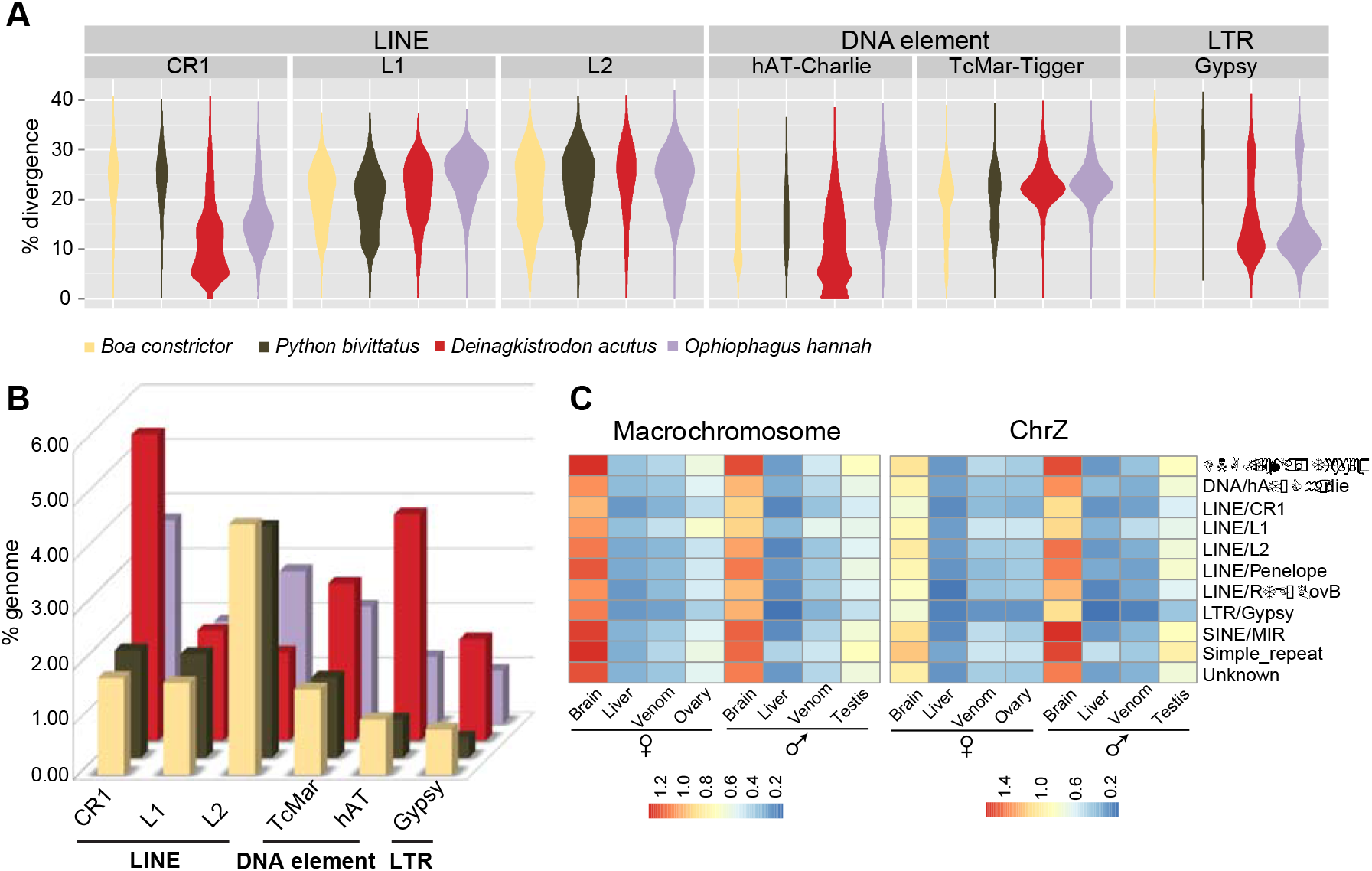
Genomic and transcriptomic variation of snake transposable elements. **(A)** Violin plots showing each type of TE’s frequency distribution of sequence divergence level from the inferred ancestral consensus sequences. Clustering of TEs with similar divergence levels, manifested as the ‘bout’ of the violin, corresponds to the burst of TE amplification. **(B)** Bar plots comparing the genome-wide TE content between four snake species. TE families were annotated combining information of sequence homology and *de novo* prediction. **(C)** TE’s average normalized expression level (measured by RPKM) across different tissues in five-pace viper.

These TEs are presumably silenced through epigenetic mechanisms to prevent their deleterious effects of transposition or mediating genomic rearrangements. Indeed, very few are transcribed in all of the tissue examined, except, unexpectedly in brain (Figure 2C). This brain-specific expression prompted us to test whether some snake TE families might have been co-opted into brain gene regulatory networks. Focusing on highly expressed (RPKM>5) TEs’ that are located within 5kb flanking regions of genes, we found that these nearby genes also show a significantly higher expression in brain than any other tissues (*P-value*<1.1e-40, Wilcoxon test, **Supplementary Figure 3**). The expression levels of individual genes are strongly correlated (*P-value*<1.35e-08, Spearman’s test) with those of nearby TE’s. These genes are predominantly enriched (*Q-value* < 0.05, Fisher’s Chi-square test, **Supplementary Data 2**) in functional domains of ‘biological process’ compared to ‘cellular component’ and ‘molecular function’, and particularly enriched categories include environmental response (‘response to organic substance’, ‘regulation of response to stimulus’ and ‘sensory perception of light stimulus’) and brain signaling pathways (‘neuropeptide signaling pathway’, ‘opioid receptor signaling pathway’ and ‘regulation of cell communication’ etc.). Further experimental studies are required to elucidate how some of these TEs evolved to regulate the brain gene expression; these results nevertheless highlighted the evolution dynamics and potential functional contribution of TEs in shaping the snake genome evolution.

## Evolution of snake genes and gene families

To pinpoint the critical genetic changes underlying the phenotypic innovations of snakes, we next mapped protein coding genes’ gain and loss (Figure 1B), signatures of adaptive or degenerative evolution (Figure 3A) measured by their ratios (*ω*) of nonsynonymous vs. synonymous substitution rates (**Supplementary Data 3**) onto the phylogenetic tree. We inferred a total of 1,725 gene family expansion and 3,320 contraction events, and identified 610 genes that appear to have undergone positive selection (PSG) and 6149 with relaxed selective constraints (RSG) at different branches, using a likelihood model and conserved lineage-specific test^28^. Genes of either scenario were separated for analyzing their enriched gene ontology (GO) and mouse orthologs’ mutant phenotype (MP) terms, assuming most of them have a similar function in snakes.

**Figure 3.**
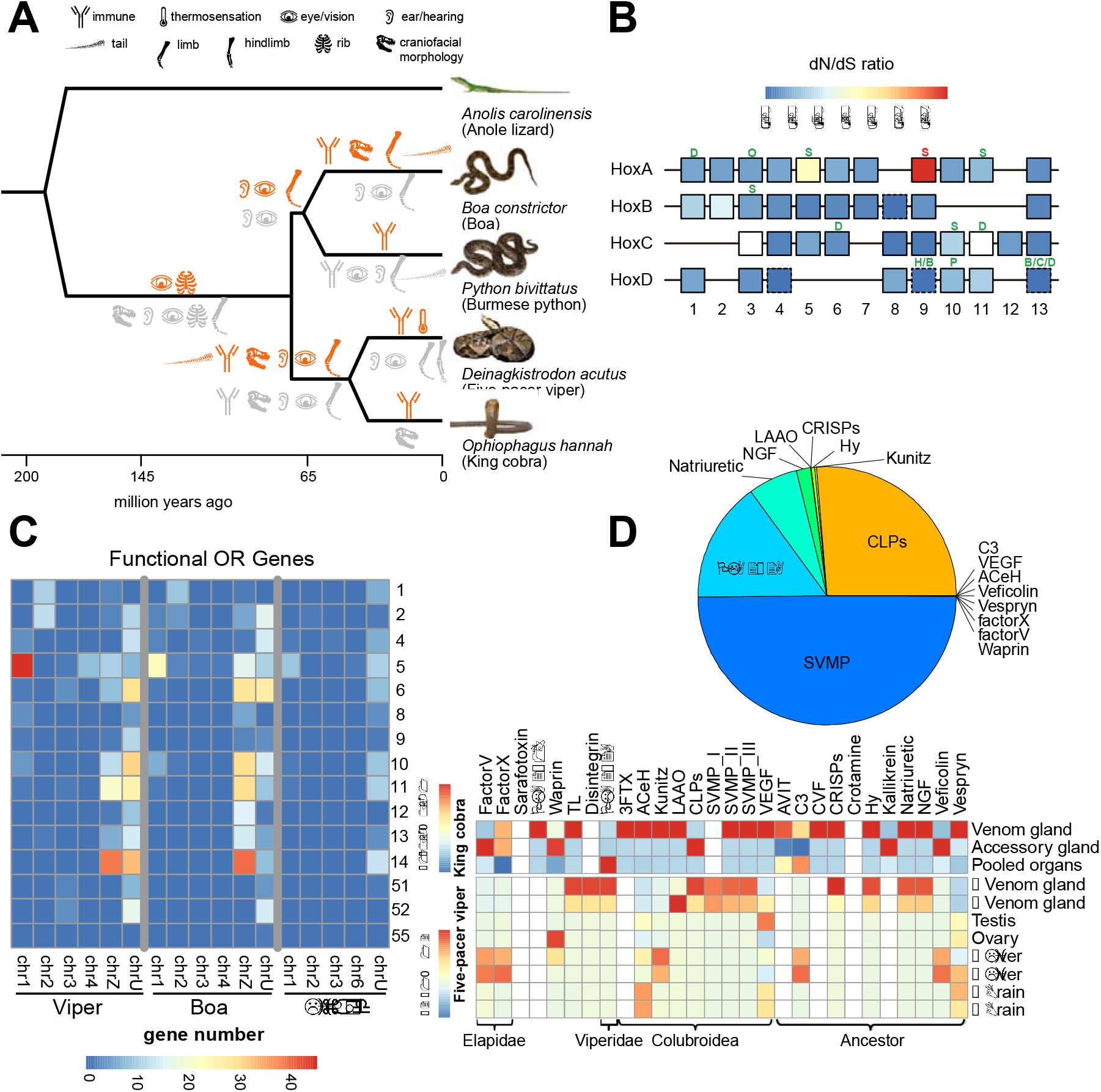
Evolution of snake genes and gene families. **(A)** Phylogenetic distribution of mutant phenotypes (MP) of mouse orthologs of snakes. Each MP term is shown by an organ icon, and significantly enriched for snake genes undergoing positive selection (red) or relaxed selective constraints (gray) inferred by lineage-specific PAML analyses. **(B)** We show the four *Hox* gene clusters of snakes, with each box showing the ratio of nonsynonymous (dN) over synonymous substitution (dS) rate at the snake ancestor lineage. White boxes represent genes that haven’t been calculated for their ratios due to the genome assembly issue in species other than five-pacer viper. Boxes with dotted line refer to genes with dS approaching 0, therefore the dN/dS ratio cannot be directly shown. Each cluster contains up to 13 *Hox* genes with some of them lost during evolution. We also marked certain *Hox* genes undergoing positive selection (in red) or relaxed selective constraints (in green) at a specific lineage above the box. Each lineage was denoted as: S: *Serpentes* (ancestor of all snakes), H: *Henophidia* (ancestor of boa and python), B: *Boa constrictor*, P: *Python bivittatus*, C: *Colubroidea,* D: *Deinagkistrodon acutus,* O: *Ophiophagus hannah.* **(C)** Comparing olfactory receptor (OR) gene repertoire between boa, viper and lizard. Each cell corresponds to a certain OR family (shown at y-axis) gene number on a certain chromosome (x-axis). **(D)** Pie chart shows the composition of normalized venom gland transcripts of male five-pacer viper. The heatmap shows the normalized expression level (in RPKM) across different tissues of viper and king cobra. We grouped the venom genes by their time of origination, shown at the bottom x-axis.

Significantly (*P-value*<0.05, Fisher’s exact test) enriched MP terms integrated with their branch information illuminated the molecular evolution history of snake-specific traits (Figure 3A). For example, as adaptations to a fossorial lifestyle, the four-legged snake ancestor^29^ had evolved an extreme elongated body plan without limbs, and also fused eyelids (‘spectacles’, presumably for protecting eyes against soil^30^). The latter is supported by the results for the PSG genes *Ereg*, and RSGs *Cecr2* and *Ext1* at the snake ancestor branch (**Supplementary Data 4**), whose mouse mutant phenotype is shown as prematurely opened or absent eyelids. The limbless body plan has already driven many comparisons of expression domains and coding-sequences of the responsible *Hox* genes between snakes and other vertebrates^7,17^. We here refined the analyses to within-snake lineages, focusing on sequence evolution of *Hox* and other genes involved in limb development and somitogenesis. We annotated the nearly complete sequences of 39 *Hox* genes organized in four clusters (*HoxA-HoxD*) of the five-pacer viper. Compared to the green anole lizard, the four studied snake species have *Hox* genes whose sizes are generally reduced, due to the specific accumulation of DNA transposons in the lizard’s introns and intergenic regions (**Supplementary Figure 4**). However, snakes have accumulated particularly higher proportions of simple tandem repeat and short interspersed element (SINE) sequences within *Hox* clusters (**Supplementary Figure 5**), either as a result of relaxed selective constraints and/or evolution of novel regulatory elements. We identified 11 *Hox* genes as RSGs and one (*Hoxa9*) as PSG (Figure 3B). Their combined information of gene function and affected snake lineage informed the stepwise evolution of snake body plan. In particular, *Hoxa5*^31^, *Hoxa11*^32^ and *Tbx5*^33^ which specifically pattern the forelimbs in mouse, have been identified as RSGs in the common ancestor of all four snakes. While *Hoxc11* and *Tbx4^34^*, which pattern the hindlimbs in mouse, and many other limb-patterning genes (e.g. *Gli3, Tbx18, Alx4*) were identified as RSGs that evolved independently on snake external branches (Figure 3B, **Supplementary Data 4**). These results provide robust molecular evidence supporting the independent loss of hindlimbs after the complete loss of forelimbs in snake ancestors. In the snake ancestor branch, we also identified RSGs *Hoxa11, Hoxc10* and *Lfng,* which are respectively associated with sacral formation^35^, rib formation^8^ and somitogenesis speed^36^ in vertebrates. Their changed amino acids and the expression domains that have expanded in snakes relative to lizards^17,19^ might have together contributed to the ‘de-regionalization’^17^ and elongation of the snake body plan. In several external branches, we identified *Hoxd13* independently as RSG. Besides its critical roles in limb/digit patterning^37^, it is also associated with termination of somitogenesis signal and is specifically silenced at the snake tail relative to lizard^7^. This suggests that body elongation may have evolved more than once among snake lineages. Overall, limb/digit/tail development MP terms are significantly enriched in RSGs at both ancestral and external branches of snakes (Figure 3A), and we identified many such genes at different snake lineages for future targeted experimental studies (**Supplementary Table 9, Supplementary Data 4**).

Another important adaption to snakes’ ancestrally fossorial and later ground surface lifestyle is the shift of their dominant source of environmental sensing from visual/auditory to thermal/chemical cues. Unlike most other amniotes, extant snake species do not have external ears, and some basal species (e.g., blindsnake) have completely lost their eyes. Consistently, we found MP terms associated with hearing/ear and vision/eye phenotypes (e.g., abnormal ear morphology, abnormal vision, abnormal cone electrophysiology) are enriched among RSGs along all major branches of snakes starting from their common ancestor (Figure 3A, **Supplementary Figure 6, Supplementary Data 4**). Gene families that have contracted in the ancestor of four studied snake species, and specifically in the viper, are also significantly (*Q-value*<9.08e-4, Fisher’s Exact Test) enriched in gene ontologies (GO) of ‘sensory perception of light stimulus (GO:0050953)’ or ‘phototransduction (GO:0007602)’ (Figure 1C, **Supplementary Data 5**). In particular, only three (*RH1, LWS* and *SWS1*) out of 13 opsin genes’ complete sequences can be identified in the viper genome, consistent with the results found in python and cobra^16^. By contrast, infrared receptor gene *TRPA1*^5^ and ubiquitous taste-signaling gene *TRPM5*^38^ have respectively undergone adaptive evolution in five-pacer viper and the ancestor of boa and python. Gene families annotated with the GO term ‘olfactory receptor (OR) activity’ have a significant (*Q-value* < 1.63e-4, Fisher’s Exact Test) expansion in all snake species studied and at some of their ancestral nodes, except for the king cobra (**Supplementary Data 6**). In the boa and viper, whose genome sequences have much better quality than the other two snake genomes, we respectively annotated 369 and 412 putatively functional OR genes, based on homology search and the characteristic 7-TM (transmembrane) structure (Materials and Methods). Both terrestrial species have an OR repertoire predominantly comprised of class II OR families (OR1-14, presumably for binding airborne molecules, Figure 3C), and their numbers are much higher than the reported numbers in other Squamate genomes^39^. Some (ranging from 18 to 24) class I (OR51-56, for water-borne molecules) genes have also been found in the two species, indicating this OR class is not unique to python as previously suggested^39^. Compared with the green anole lizard, the boa and viper exhibit a significant (*P*<0.05, Fisher’s exact test) size expansion of OR family 5, 11 and 14, and also a bias towards being located on the Z chromosome (Figure 3C), leading to higher expression of many OR genes in males than in females (see below). In particular, OR5 in the viper probably has experienced additional expansion events and become the most abundant (with 71 members) family in the genome. Intriguingly, this family is specifically enriched in birds of prey^40^ relative to other birds, and in non-frugivorous bats vs. frugivorous bats^41^. Therefore, its expansion in the five-pacer viper could have been positively selected for a more efficient detection of prey.

Besides acute environmental sensing, specialized fangs^6^ and venoms^11^ (e.g, hemotoxins of viper or neurotoxins of elapid) arm the venomous snakes (~650 species) to immediately immobilize the much larger prey for prolonged ingestion, which probably comprised one of the most critical factors that had led to the advanced snakes’ species radiation. It has been proposed that the tremendous venom diversity probably reflects snakes’ local adaption for the prey^42^ and was generated by changes in the expression of pre-existing or duplicated genes^11,43^. Indeed, we found that the five-pacer viper’s venom gland gene repertoire has a very different composition comparing to other viper^44^ or elapid species^10^ (Figure 3D). We have annotated a total of 35 venom genes or gene families using all the known snake venom proteins as the query. Certain gene families, including snake venom metalloproteinases (SVMP), C-type lectin-like proteins (CLPs), thrombin-like snake venom serine proteinases (TL), Kunitz and disintegrins, have more genomic copies in the five-pacer viper than other studied snakes or the green anole lizard (**Supplementary Table 10**); while characteristic elapid venom genes like three-finger toxins (3FTx) are absent from the viper genome. Most venom proteins of both the viper and king cobra have expression restricted to venom or accessary glands, and for both species this is particularly seen for those genes that originated in the ancestor of snakes or of advanced snakes (Figure 3D). But for elapid or viper specific venom genes, i.e., those that originated more recently, they usually express in the liver of the other species. Such cases include FactorV, FactorX of king cobra and PLA2-2A of viper (Figure 3D). This suggests that these venom genes may have originated from metabolic proteins and undergone neo-/sub-functionalization, with altered expression.

## Evolution of snake sex chromosomes

Different snake species exhibit a continuum of sex chromosome differentiation: pythons and boas possess homomorphic sex chromosomes, which is assumed to be the ancestral state; the lack of differentiation between the W and Z chromosomes suggests that most regions of this chromosome pair recombine like the autosomes^45^. Advanced snakes usually have heteromorphic sex chromosomes that have undergone additional recombination suppressions^45,46^. We found the five-pacer viper probably has suppressed recombination throughout almost the entire sex chromosome pair, as the read coverage in the female that we sequenced is half that in the male (Figure 1C, Figure 4). By contrast, boa’s homologous chromosomal regions show a read coverage pattern that does not differ from that of autosomes. Assuming that these two species share the same ancestral snake sex-determining region, this suggests that that region is not included in our current chromosomal assembly.

**Figure 4.**
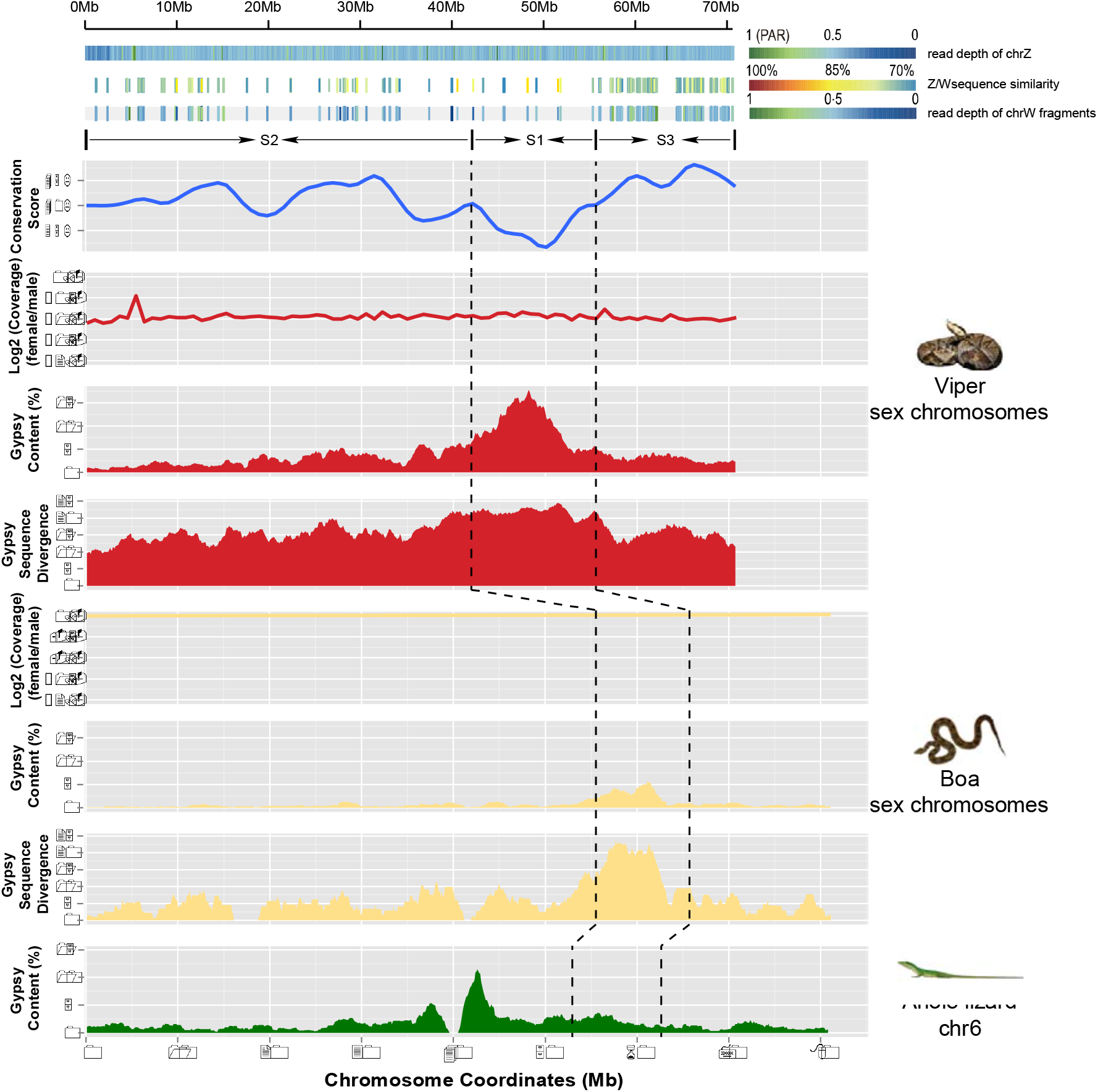
Snake sex chromosomes have at least three evolution strata. The three tracks in the top panel shows female read depths along the Z chromosome relative to the median depth value of autosomes, Z/W pairwise sequence divergence within intergenic regions, and female read depths of W-linked sequence fragments relative to the median depth value of autosomes. Depths close to 1 suggest that the region is a recombining pseudoautosomal region (PAR), whereas depths of 0.5 are expected in a highly differentiated fully sex-linked region where females are hemizygous. The identifiable W-linked fragments are much denser at the region 56Mb~70Mb, probably because this region (denoted as stratum 3, S3) has suppressed recombination most recently. S2 and S1 were identified and demarcated by characterizing the sequence conservation level (measured by LASTZ alignment score, blue line) between the chrZs of boa and viper. At the oldest stratum S1 where recombination has been suppressed for the longest time, there is an enrichment of repetitive elements on the affected Z-linked region (Gypsy track in red, 100kb non-overlapping sliding window). And these Z-linked TEs A similar pattern was found in homologous recombining region of boa, but not in lizard.

In plants, birds and mammals, it has been found that recombination suppression probably occurred by a succession of events. This has led to the punctuated accumulation of excessive neutral or deleterious mutations on the Y or W chromosome by genetic drift, and produced a gradient of sequence divergence levels over time, which are termed ‘evolutionary strata’^47-49^. Advanced snakes have been suggested to have at least two strata^12^. One goal of our much better genome assembly of the five-pacer viper compared to those of any other studied advanced snakes^10,12^ (**Supplementary Table 4**) was to reconstruct a fine history of snake sex chromosome evolution. We assembled 77Mb Z-linked and 33Mb W-linked scaffolds (**Materials and Methods**). The reduction of female read coverage along the Z chromosome suggests there is a substantial divergence between Z- and W- linked sequences, and this would enable the assembly of two chromosomes’ scaffolds in separate. Mapping the male reads confirmed that the inferred W-linked scaffold sequences are only present in female (**Supplementary Figure 7**). Their density and pairwise sequence divergence values within putative neutral regions along the Z chromosome indicate at least two ‘evolution strata’, with the older stratum extending 0~56Mb, and the younger one extending 56~70Mb. The boundary at 56Mb region can be also confirmed by analyses of repetitive elements on the Z chromosome (see below). Consistently, identifiable W-linked fragments are found at the highest density per megabase in the latter (Figure 4), suggesting that this region has suppressed recombination more recently. The older stratum includes much fewer identifiable fragments that can resolve the actual times of recombination suppression events. To study this region further, we inspected the homologous Z-linked region, whose recombination has also been reduced, albeit to a much smaller degree than the W chromosome, after the complete suppression of recombination between Z/W in females. In addition, Z chromosome is biasedly transmitted in males. As males usually have a higher mutation rate than females, due to many more rounds of DNA replication during spermatogenesis than oogenesis (‘male-driven evolution’)^50^, Z-linked regions are expected to have a generally higher mutation rate than any other regions in the genome. This male-driven evolution effect has been demonstrated in other snake species^12^, and also been validated for the snakes inspected in this study (**Supplementary Figure 8**). As a result, we expected regions in older strata should be more diverged from their boa autosome-like homologs than those in the younger strata. This enabled us to identify another stratum (0~42Mb, stratum 2, S2 in Figure 4) and demarcate the oldest one (42~56Mb, S1), by estimating the sequence conservation level (measured by LASTZ alignment score, blue line) between the Z chromosomes of boa and viper. The Z-linked region in the inferred oldest stratum S1 exhibits the highest sequence divergence with the homologous W-linked region, and also the highest proportion of repetitive elements (CR1, Gypsy and L1 elements; Figure 4 shows the example of Gypsy; other repeats are shown in **Supplementary Figure 9**). This can be explained by the effect of genetic drift^51^, which has been acting on the Z-linked S1 for the longest time since it reduced recombination rate in females. As a result, the accumulated repeats of S1 also tend to have a higher divergence level from the inferred ancestral consensus sequences compared to nearby strata (Figure 4). Unexpectedly, a similar enrichment was found in the homologous region of S1 in boa, despite being a recombining region and exhibiting the same coverage depth between sexes (Figure 4, **Supplementary Figure 9**). This indicates the pattern is partially contributed by the ancestral repeats that had already accumulated on the proto-sex chromosomes of snake species. Since our current viper sex chromosomal sequences used the green anole lizard chromosome 6 as a reference, rearrangements within this chromosome make it impossible to test whether S2 encompasses more than one stratum.

We dated the three resolved strata by constructing phylogenetic trees with homologous Z- and W- linked gene sequences of multiple snake species. Combining the published CDS sequences of pygmy rattlesnake (Viperidae family species) and garter snake (Colubridae family species)^12^, we found 31 homologous Z-W gene pairs, representing the three strata. All of them clustered by chromosome (i.e., the Z-linked sequences from all the species cluster together, separately from the W-linked ones) rather than by species (**Supplementary Figure 10-12**). This indicates that all three strata formed before the divergence of advanced snakes, and after their divergence from boa and python, i.e., about 66.9 million years ago (Figure 1C).

We found robust evidence of functional degeneration on the W chromosome. It is more susceptible to the invasion of TEs: the assembled sequences’ overall repeat content is at least 1.5 fold higher than that of the Z chromosome, especially in the LINE L1 (2.9 fold) and LTR Gypsy families (4.3 fold) (**Supplementary Table 11 and Supplementary Figure 13**). Of 1,135 Z-linked genes, we were only able to identify 137 W-linked homologs. Among these, 62 (45.26%) have probably become pseudogenes due to nonsense mutations (**Supplementary Table 12**). W-linked loci generally transcribe at a significantly (*P-value*<0.0005, Wilcoxon test) lower level, with pseudogenes transcribing even lower relative to the autosomal or Z-linked loci regardless of the tissue type (**Supplementary Figure 14-15**). Given such a chromosome-wide gene loss, like other snakes^12^ or majority of species with ZW sex chromosomes^52^, the five-pacer viper shows a generally male-biased gene expression throughout the Z-chromosome and probably has not evolved global dosage compensation (**Supplementary Figure 16**).

## Discussion

The ‘snake-like’ body plan has evolved repeatedly in other tetrapods (e.g., worm lizard and caecilians), in which limb reduction/loss seems to have always been accompanied by the body elongation. For example, several limb-patterning *Hox* genes (*Hoxc10*, *Hoxd13*) identified as RSG also have been characterized by previous work with a changed expression domain along the snake body axis^7,17^. Another RSG *Hoxa5* which is involved in the forelimb patterning^31^, also participates in lung morphogenesis^53^. It might have been involved in the elimination of one of the snake lungs during evolution. Therefore, the newly identified PSGs and RSGs throughout the snake phylogeny in this work (**Supplementary Data 3-4, Supplementary Table 9**) can provide informative clues for future experimental work to use snake as an emerging ‘evo-devo’ model^54^ to understand the genomic architecture of developmental regulatory network of organogenesis, or the crosstalk between these networks.

Like many of its other reptile relatives, the snake ancestor is very likely to determine the sex by temperature and does not have sex chromosomes. Extant species Boa can still undergo occasional parthenogenesis and is able to produce viable WW offspring^55^, consistent with its the most primitive vertebrate sex chromosome pair reported to date. In the ancestor of advanced snakes, we inferred that there were at least three recombination suppressions occurred between Z/W, leading to a generally degenerated W chromosome that we have observed in five-pacer viper. How snakes genetically determine the sex would be a rather intriguing question to study in future.

## Materials and Methods

### Genome sequencing and assembly

All animal procedures were carried out with the approval of China National Genebank animal ethics committee. We extracted genomic DNAs from blood of a male and a female five-pacer viper separately. A total of 13 libraries with insert sizes ranging from 250bp to 40kb were constructed using female DNA, and 3 libraries with insert sizes from 250 bp to 800 bp were constructed using male DNA. We performed paired-end sequencing (HiSeq 2000 platform) following manufacturer’s protocol, and produced 528 Gb raw data (357 Gb for female and 171 Gb for male). We estimated the genome size based on the K-mer distribution. A K-mer refers to an artificial sequence division of K nucleotides iteratively from sequencing reads. The genome size can then be estimated through the equation G=K_num/Peak_depth, where the K_num is the total number of K-mer, and Peak_depth is the expected value of K-mer depth^56^. We found a single main peak in the male K-mer (K=17) frequency distribution and an additional minor peak in the female data, the latter of which probably results from the divergence between W and Z chromosomes (**Supplementary Figure 1**). Based on the distribution, we estimated that the genome size of this species is about 1.43 Gb (**Supplementary Table 2**), comparable to that of other snakes (1.44 Gb and 1.66Gb for Burmese python and King cobra^10,16^, respectively).

After filtering out low-quality and duplicated reads, we performed additional filtering using the following criteria: we excluded the reads from short-insert libraries (250, 500, 800 bp) with ‘N’s over 10% of the length or having more than 40 bases with the quality lower than 7, and the reads from large-insert libraries (2 kb to 40 kb) with ‘N’s over 20% of the length or having more than 30 bases with the quality lower than 7. Finally, 109.20 Gb (73X coverage) male reads and 148.49Gb (99X coverage) female reads were retained for genome assembly (Supplementary Table 1) using SOAPdenovo^57^ (http://soap.genomics.org.cn). To assemble the female and male genomes, reads from small-insert libraries of the female and male individual were used for contig construction separately. Then read-pairs from small- and large-insert libraries were utilized to join the contigs into scaffolds. We also used female long-insert libraries to join the male contigs into the longer scaffolds. At last, small-insert libraries of female and male individuals were used for gap closure for their respective genomes. The final assemblies of female and male have a scaffold N50 length of 2.0 Mb and 2.1 Mb respectively, and the gap content of the two genomes are both less than 6% (♀ 5.29%, ♂ 5.61%)(**Supplementary Table 4**).

To access the assembly quality, reads from small-insert libraries that passed our filtering criteria were aligned onto the two assemblies using BWA^58^ (Version: 0.5.9-r16) allowing 8 mismatches and 1 indel per read. A total of ~97% reads can be mapped back to the draft genome (**Supplementary Table 3**), spanning 98% of the assembled regions excluding gaps (**Supplementary Table 13**), and most genomic bases were covered by about 80X reads (**Supplementary Figure 17**). Thus we conclude that we have assembled most part of the five-pacer viper genome. To further test for potential mis-joining of the contigs into scaffolds, we analyzed the paired-end information and found that 57% of the paired-end reads can be aligned uniquely with the expected orientation and distance. This proportion of the long insert library is significantly lower than that from the short insert libraries due to a circulization step during the library construction. When such paired-ends were excluded, the proportion increased to 94.98% (**Supplementary Table 3**). Overall, these tests suggested that the contigs and scaffolds are consistent with the extremely high density of paired-end reads, which in turn indicated the high-quality of the assembly.

Previous cytogenetic studies showed that snake genomes show extensive inter-chromosomal conservation with lizard^27,45^. Thus we used the chromosomal information from green anole lizard^24^ as a proxy to assign the snake scaffolds. We first constructed their orthologous relationship combining information of synteny and reciprocal best BLAST hits (RBH). Then gene coordinates and strandedness from the consensus chromosome were used to place and orient the snake scaffolds. Furthermore, we linked scaffolds into chromosomes with 600 ‘N’s separating the adjacent scaffolds. In total, 625 five-pacer viper scaffolds comprising 832Mb (56.50% scaffolds in length) were anchored to 5 autosomes and Z chromosome (**Supplementary Table 6**).

### Repeat and gene annotation

We identified the repetitive elements in the genome combining both homology-based and *de novo* predictions. We utilized the ‘Tetrapoda’ repeat consensus library in Repbase^59^ for RepeatMasker (http://www.repeatmasker.org) to annotate all the known repetitive elements in the five-pacer viper genome. In order to maximize the identification and classification of repeat elements, we further used RepeatModeler (http://www.repeatmasker.org/RepeatModeler.html) to construct the consensus repeat sequence libraries of the green anole lizard, boa and five-pacer viper, then used them as a query to identify repetitive elements using RepeatMasker. Finally, we retrieved a non-redundant annotation for each species after combining all the annotation results using libraries of ‘Tetrapoda’, ‘green anole lizard’, ‘boa’ and ‘five-pacer viper’. For the purpose of comparison, we ran the same pipeline and parameters in all the snake and lizard genomes as shown in **Supplementary Table 7**. To provide a baseline estimate for the sequence divergence of TEs from the snake ancestral status, we first merged the genomes from boa and five-pacer viper, and constructed the putative ancestral consensus sequences using RepeatModeler. Then TE sequences of each snake species were aligned to the consensus sequence to estimate their divergence level using RepeatMasker.

For gene annotation, we combined resources of sequence homology, *de novo* prediction and transcriptome to build consensus gene models of the reference genome. Protein sequences of green lizard, chicken and human were aligned to the reference assembly using TBLASTN (E-value <= 1E-5)^60^. Then the candidate gene regions were refined by GeneWise^61^ for more accurate splicing sites and gene models. We randomly selected 1000 homology-based genes to train Augustus^62^ for *de novo* prediction on the pre-masked genome sequences. We mapped RNA-seq reads of 13 samples to the genome using TopHat (v1.3.1)^63^ and then assembled the transcripts by Cufflinks (v1.3.0) (http://cufflinks.cbcb.umd.edu/). Transcripts from different samples were merged by Cuffmerge. Finally, gene models from these three methods were combined into a non-redundant gene set.

We finally obtained 21,194 protein-coding genes with intact open reading frames (ORFs) (**Supplementary Table 14**). The gene models (measured by gene length, mRNA length, exon number and exon length) are comparable to those of other vertebrates and are well supported by the RNA-Seq data (**Supplementary Figure 18 and Supplementary Table 5**). To annotate the gene names for each predicted protein-coding locus, we first mapped all the 21,194 genes to a manually collected Ensembl gene library, which consists of all proteins from *Anolis carolinensis, Gallus gallus, Homo sapiens, Xenopus tropicalis and Danio rerio.* Then the best hit of each snake gene was retained based on its BLAST alignment score, and the gene name of this best hit gene was assigned to the query snake gene. Most of the predicted genes can be found for their orthologous genes in the library at a threshold of 80% alignment rate (the aligned length divided by the original protein length), suggesting our annotation has a high quality (**Supplementary Table 15**).

### RNA-seq and gene expression analyses

Total RNAs were isolated from four types of tissues collected from both sexes, including brain, liver, venom gland and gonad (**Supplementary Table 16**). RNA sequencing libraries were constructed using the Illumina mRNA-Seq Prep Kit. Briefly, oligo(dT) magnetic beads were used to purify poly-A containing mRNA molecules. The mRNAs were further fragmented and randomly primed during the first strand synthesis by reverse transcription. This procedure was followed by a second-strand synthesis with DNA polymerase I to create double-stranded cDNA fragments. The cDNAs were subjected to end-repairing by Klenow and T4 DNA polymerases and A-tailed by Klenow lacking exonuclease activity. The fragments were ligated to Illumina Paired-End Sequencing adapters, size selected by gel electrophoresis and then PCR amplified to complete the library preparation. The paired-end libraries were sequenced using Illumina HiSeq 2000 (90/100 bp at each end).

We used TopHat (v1.3.1) for aligning the RNA-seq reads and predicting the splicing junctions with the following parameters: -I/--max-intron-length: 10000, --segment-length: 25, --library-type: fr-firststrand, --mate-std-dev 10, -r/--mate-inner-dist: 20. Gene expression was measured by reads per kilobase of gene per million mapped reads (RPKM). To minimize the influence of different samples, RPKMs were adjusted by a scaling method based on TMM (trimmed mean of M values; M values mean the log expression ratios)^64^ which assumes that the majority of genes are common to all samples and should not be differentially expressed.

### Evolution analyses

A phylogenetic tree of the five-pacer viper and the other sequenced genomes (*Xenopus tropicalis, Homo sapiens, Mus musculus, Gallus gallus, Chelonia mydas, Alligator mississippiensis, Anolis carolinensis, Boa constrictor, Python bivittatus* and *Ophiophagus hannah*) was constructed using the 5,353 orthologous single-copy genes. Treebest (http://treesoft.sourceforge.net/treebest.shtml) was used to construct the phylogenetic tree. To estimate the divergence times between species, for each species, 4-fold degenerate sites were extracted from each orthologous family and concatenated to one sequence for each species. The MCMCtree program implemented in the Phylogenetic Analysis by Maximum Likelihood (PAML)^65^ package was used to estimate the species divergence time. Calibration time was obtained from the TimeTree database (<u>http://www.timetree.org/</u>). Three calibration points were applied in this study as normal priors to constrain the age of the nodes described below. 61.5-100.5 MA for the most recent common ancestor (TMRCA) of human-mouse; 259.7-299.8 MA for TMRCA of Crocodylidae and Lepidosauria; 235–250.4 MA for TMRCA of Aves and Crocodylidae^66^.

To examine the evolution of gene families in Squamate reptiles, genes from four snakes (*Boa constrictor, Python bivittatus, Deinagkistrodon acutus, Ophiophagus hannah*) and green anole lizard were clusterred into gene families by Treefam (min_weight=10, min_density=0.34, and max_size=500)^67^. The family expansion or contraction analysis was performed by CAFE^68^. In CAFE, a random birth-and-death model was proposed to study gene gain and loss in gene families across a user-specified phylogenetic tree. A global parameter *λ* (lambda), which described both gene birth (λ) and death (μ = -λ) rate across all branches in the tree for all gene families was estimated using maximum likelihood method. A conditional p-value was calculated for each gene family, and the families with conditional p-values lower than 0.05 were considered to have a significantly accelerated rate of expansion and contraction.

For the PAML analyses, we first assigned orthologous relationships for 12,657 gene groups among all Squamata and outgroup (turtle) using the reciprocal best blast hit algorithm and syntenic information. We used PRANK^69^ to align the orthologous gene sequences, which takes phylogenetic information into account when placing a gap into the alignment. We filtered the PRANK alignments by gblocks^70^ and excluded genes with high proportion of low complexity or repetitive sequences to avoid alignment errors. To identify the genes that evolve under positive selection (PSGs), we performed likelihood ratio test (LRT) using the branch model by PAML^65^. We first performed a LRT of the two-ratio model, which calculates the dN/dS ratio for the lineage of interest and the background lineage, against the one-ratio model assuming a uniform dN/dS ratio across all branches, so that to determine whether the focal lineage is evolving significantly faster (p-value < 0.05). In order to differentiate between episodes of positive selection and relaxation of purifying selection (RSGs), we performed a LRT of two-ratios model against the model that fixed the focal lineage’s dN/dS ratio to be 1 (p-value < 0.05) and also required PSGs with the free-ratio model dN/dS > 1 at the focal lineage. For the identified RSGs and PSGs, we used their mouse orthologs’ mutant phenotype information^71^ and performed enrichment analyses using MamPhEA^72^. Then we grouped the enriched MP terms by different tissue types.

### Olfactory receptor (OR), *Hox* and venom gene annotation

To identify the nearly complete functional gene repertoire of OR, *Hox* and venom toxin genes in the investigated species, we first collected known amino acid sequences of 458 intact OR genes from three species (green anole lizard, chicken and zebra finch)^73^, all annotated *Hox* genes from *Mus musculus* and *HoxC3* from *Xenopus tropicalis*, and obtained the query sequences of a total of 35 venom gene families^74^ from UniProt (<u>http://www.uniprot.org/</u>) and NCBI (<u>http://www.ncbi.nlm.nih.gov/</u>). These 35 venom gene families represent the vast majority of known snake venoms. Then we performed a TBlastN^60^ search with the cutoff E-value of 1E-5 against the genomic data using these query sequences. Aligned sequence fragments were combined into one predicted gene using perl scripts if they belonged to the same query protein. Then each candidate gene region was extended for 2kb from both ends to predict its open reading frame by GeneWise^61^. Obtained sequences were verified as corresponding genes by BlastP searches against NCBI nonredundant (nr) database. Redundant annotations within overlapped genomic regions were removed.

For the OR gene prediction, these candidates were classified into functional genes and nonfunctional pseudogenes. If a sequence contained any disruptive frame-shift mutations and/or premature stop codons, it was annotated as a pseudogene. The remaining genes were examined using TMHMM2.0^75^. Those OR genes containing more at least 6 transmembrane (TM) structures were considered as intact candidates and the rest were also considered as pseudogenes. Finally, each OR sequence identified was searched against the HORDE (the Human Olfactory Data Explorer) database (http://genome.weizmann.ac.il/horde/) using the FASTA (ftp://ftp.virginia.edu/pub/fasta) and classified into the different families according to their best-aligned human OR sequence. For the venom toxin genes, we only kept these genes with RPKM higher than 1 in the five-pacer viper and king cobra venom gland tissue as final toxin gene set.

### Identification and analyses of sex-linked genes

To identify the Z-linked scaffolds in the male assembly, we aligned the female and male reads to the male genome separately with BWA^58^ allowing 2 mismatches and 1 indel. Scaffolds with less than 80% alignment coverage (excluding gaps) or shorter than 500 bp in length were excluded. Then single-base depths were calculated using SAMtools^76^, with which we calculated the coverage and mean depth for each scaffold. The expected male vs. female (M:F) scaled ratio of a Z-linked scaffold is equal to 2, and we defined a Z-linked scaffold with the variation of an observed scaled ratio to be less than 20% (i.e. 1.6 to 2.4). With this criteria, we identified 139 Z-lined scaffolds, representing 76.93Mb with a scaffold N50 of 962 kb (**Supplementary Table 17**). These Z-linked scaffolds were organized into pseudo-chromosome sequence based on their homology with green anole lizard. Another characteristic pattern of the Z-linked scaffolds is that there should be more heterozygous SNPs in the male individual than in the female individual resulted from their hemizygous state in female. We used SAMtools^76^ for SNP/indel calling. SNPs and indels whose read depths were too low (<10) or too high (>120), or qualities lower than 100 were excluded. As expected, the frequency of heterozygous sites of Z chromosome of the female individual is much lower than that of the male individual (0.005% vs 0.08%), while the heterozygous rate of autosomes are similar in both sex (~0.1%) (**Supplementary Table 18**). To identify the W-linked scaffolds, we used the similar strategy as the Z-linked scaffold detection to obtain the coverage and mean depth of each scaffold. Then we identified those scaffolds covered by female reads over 80% of the length, and by male reads with less than 20% of the length. With this method, we identified 33 Mb W-linked scaffolds with a scaffold N50 of 48 kb (**Supplementary Table 19**).

We used the protein sequences of Z/W gametologs from garter snake and pygmy rattle snake^12^ as queries and aligned them to the genomes of boa (the SGA assembly, http://gigadb.org/dataset/100060), five-pacer viper and king cobra with BLAST^60^. The best aligned (cutoff: identity>=70%, coverage>=50%) region with extended flanking sequences of 5kb at both ends was then used to determine whether it contains an intact open reading frame (ORF) by GeneWise^61^ (-tfor -genesf -gff -sum). We annotated the ORF as disrupted when GeneWise reported at least one premature stop codon or frame-shift mutation. CDS sequences of single-copy genes’ Z/W gametologs were aligned by MUSCLE^77^ and the resulting alignments were cleaned by gblocks^70^ (-b4=5, -t=c, -e=-gb). Only alignments longer than 300bp were used for constructing maximum likelihood trees by RAxML^78^ to infer whether their residing evolutionary stratum is shared among species or specific to lineages.

## Supplementary Figure Legend

**Supplementary Figure 1.** K-mer estimation of the genome size of five-pacer viper. Distribution of 17-mer frequency in the used sequencing reads from female (left) and male (right) samples. The x-axis represents the sequencing depth. The Y-axis represents the proportion of a K-mer counts in total K-mer counts at a given sequencing depth. The estimated genome size is about 1.43 Gb.

**Supplementary Figure 2.** GC isochore structure of different tetrapod genomes. We show the standard deviation (SD) of GC content calculated with different window size (3 kb to 320 kb) for different vertebrate genomes.

**Supplementary Figure 3.** Comparing expression levels of genes nearby expressed TEs in each tissue.

We show expression patterns of genes around highly expressed TE (RPKM > 5) in different tissues from both sexes, including brain, liver, venom gland and gonad. We also performed comparison between the focal tissue vs. the other tissues. We show levels of significance with asterisks. *: 0.001=< *P*-value <0.01; **: 0.0001 =< *P*<0.001; ***: *P*<0.0001.

**Supplementary Figure 4.** Comparison of *Hox* gene structure between snakes and lizard. Schematic representation of four *Hox* clusters in anole lizard, boa, Burmese python, five-pacer viper and king cobra. Each number from 1 to 13 denotes the specific *Hox* gene belonging to one cluster. We showed the length difference between each species vs. mouse by the colored lines (for intergenic regions) or boxes (for intronic regions): a 1.5~3 fold increase of length was shown by blue, a more than 3-fold increase was shown in red. Exons were shown by vertical lines, and dotted lines refer to exons with unknown boundaries, either due to assembly issues. Double-slashes refer to the gap between two different scaffolds.

**Supplementary Figure 5.** Repeat accumulation at *Hox* gene clusters.

Comparison of the TE and simple repeat content of *Hox* cluster genes with 5kb flanking regions between snakes and lizard. We calculated the repeat density by dividing the total length of specific repeat sequence vs. the length of corresponding region. This density was normalized over the genome-wide repeat density and then shown by heatmap.

**Supplementary Figure 6.** Phylogenetic distribution of enriched MP terms.

We identified enriched mutant phenotypes (MP) of mouse orthologs of snake genes that are undergoing lineage-specific positive selection (red) and relaxed selective constraints (gray). And then we mapped these MP terms onto the snake phylogeny.

**Supplementary Figure 7.** Read coverage density plot of different linkage groups.

For each linkage group (from left to right, chrW, chrZ, chr1), male reads were plotted in blue, and female reads in red. The identified chrW scaffolds in this work all show a female-specific read depth pattern.

**Supplementary Figure 8.** Male-driven evolution effect in snakes.

For each gene, we calculated the substitution rates between anole lizard and each of boa, Burmese python, five-pacer viper and king cobra at synonymous sites (dS) and non-synonymous sites (dN) divided into different chromosome sets. To detect branch-specific differences, we obtained for each gene the ratios of these evolutionary rates between the different snake species vs. boa. Since boa’s homologous chromosomal region to the Z chromosomes of other advanced snakes represent the ancestral status of snake sex chromosomes, a higher relative ratios of Z-linked dS of advanced snakes vs. boa than those of autosomes indicate the male-driven evolution effect. We shown the Wilcoxon test significant differences between the Z chromosome and the autosomes (chr1-5) are marked with asterisks (***, *P*-value < 0.001).

**Supplementary Figure 9.** Repeat accumulation along the snake sex chromosomes. Shown are comparisons of the distribution of Gypsy, CR1 and L1 content along the chromosome in anole lizard, Boa and five-pacer viper. The TE content was calculated by averaging the TE density of each sliding window of 100kb as well as the flanking 10 windows.

**Supplementary Figure 10.** Gene trees for Z and W linked gametologs in S1 region. Shown are maximum likelihood (ML) trees using coding regions of Z and W allelic sequences from multiple snake species, with the gene name under each tree and bootstrapping values at each node. Trees that show separate clustering of Z- or W- linked gametologs provide strong evidence that these genes suppressed recombination before the speciation.

**Supplementary Figure 11.** Gene trees for Z and W linked gametologs in S2 region.

**Supplementary Figure 12.** Gene trees for Z and W linked gametologs in S3 region.

**Supplementary Figure 13.** Comparison of repeat content between viper chrZ and chrW. TE families were determined based on the combined annotations of Repbase, RepeatModeler, and coverage in the genome was annotated using in-house scripts.

**Supplementary Figure 14.** Comparison of gene expression across different chromosomes.

We show gene expression patterns between different chromosome sets across tissues. ‘pseudo-W’ refers to W-linked genes that have premature stop codons or frameshift mutations

**Supplementary Figure 15.** Pairwise comparison of gene expression levels between homologous Z and W alleles.

**Supplementary Figure 16.** Gene expression along the Z chromosome and autosome chr5 in different tissues of five-pacer viper.

We show log-based male-to-female gene expression ratio along the Z chromosomes and autosome chr5. Only genes with RPKM >= 1 in both the male and female were considered. If genes are mostly non-biased, the line is expected to centered at 0. The pattern indicates five-pacer viper lacks chromosome-wide dosage compensation.

**Supplementary Figure 17.** Frequency distribution of sequencing depth.

Distribution of sequencing depth of the assembled female (left) and male (right) genomes by reads from the female and male samples. The peak depth is 76X and 77X for the female and male reads aligned to corresponding assembly, respectively.

**Supplementary Figure 18.** Comparisons of gene parameters among the sequenced representative species.

We used the published genomes of *Gallus gallus, Homo sapiens, Anolis carolinensis, Boa constrictor* to compare with *Deinagkistrodon acutus*, without finding any obvious differences between them and five-pacer viper in the annotated genes’ length and number. This indicates the high quality of gene annotation.

## Supplementary Table Legend

**Supplementary Table 1.** Statistics of five-pacer viper genome sequencing. Supplementary Table 2. Statistics of 17-mer analysis.

**Supplementary Table 3.** Statistics of reads of small-insert and large-insert libraries aligned to the male assembly.

*PE mapped* refer to reads being mapped to the genome as read pairs, and *SE mapped* represent reads being mapped to the genome as single reads.

**Supplementary Table 4.** Summary of the five-pacer viper genome assemblies.

**Supplementary Table 5.** Number of expressed genes of five-pacer viper.

**Supplementary Table 6.** Number of genes and scaffold size organized into chromosomes.

**Supplementary Table 7.** Comparison of repeat content between snakes and lizard.

**Supplementary Table 8.** Comparison of genome assembly quality between snakes and lizard.

**Supplementary Table 9.** Evolution of candidate limb-patterning genes in snakes. Candidate limb-patterning genes were collected from MGI database and published paper. Genes undergoing positive selection (P) or relaxed selective constraints (R), were identified by PAML analyses. N: there is no significant selection signal. ‘-’ refers to genes that cannot be completely assembled for their coding sequences due to genome assembly gaps.

**Supplementary Table 10.** Comparison of venom genes between snakes.

Statistics of venom gene families in the four snakes and anole lizard genomes based on homology-based prediction. AVIT: Prokineticin; C3: complement C3; CVF: Cobra Venom Factor; CRISPs: Cysteine-Rich Secretory Proteins; Hy: Hyaluronidases; Natriuretic: Natriuretic peptide; NGF: Snake Venom Nerve Growth Factors; PLA2-2A: Snake Venom Phospholipase A2 (type IIA); SVMP: Snake Venom Metalloproteinases; TL: thrombin-like snake venom serine proteinases; LAAO: Snake Venom L-Amino Acid Oxidases; PDE: phosphodiesterases; CLPs: snake C-type lectin-like proteins; VEGF: vascular endothelin growth factor; PLA2-1B: Snake Venom Phospholipase A2 (type IB); 3FTX: The three-finger toxins; ACeH: Acetylcholinesterase;

**Supplementary Table 11.** Comparison of repeat content between snake sex chromosomes and their lizard homolog.

**Supplementary Table 12.** Location of W-linked putative pseudogenes.

**Supplementary Table 13.** Fractions of bases covered by reads in the male assembly.

**Supplementary Table 14.** Characteristics of predicted protein-coding genes in the male assembly.

**Supplementary Table 15.** Number of predicted genes that can find homologs in the Ensembl library with different aligning rate cutoff.

Alignment rate was calculated by dividing the aligned length vs. the original protein length. And we required both the query and subject to satisfy our alignment cutoff. The Ensembl library consists of all proteins from Anolis carolinensis, Gallus gallus, Homo sapiens, Xenopus tropicalis and Danio rerio.

**Supplementary Table 16.** Data production and alignment statistic of RNA-Seq aligned to male genome assembly.

**Supplementary Table 17.** Statistics of the identified Z-linked scaffolds.

**Supplementary Table 18.** Statistics of SNPs identified in the female and male individual.

**Supplementary Table 19.** Statistics of identified W-linked scaffolds.

## Supplementary Data Legend

**Supplementary Data 1**

Comparing five-pacer viper’s chromosomal assignment vs. reported fluorescence in situ hybridization results.

**Supplementary Data 2**

GO enrichment of nearby genes of TE highly expressed in brain.

**Supplementary Data 3**

Positively selected genes (PSGs) and genes with relaxed selective constraints (RSGs) and their affected lineage.

**Supplementary Data 4**

Mouse mutant terms’ enrichment analyses of PSGs and RSGs across different snake branches.

**Supplementary Data 5**

GO enrichment of expanded and contracted gene families across different snake lineages.

**Supplementary Data 6**

Plots of GO enrichment of expanded and contracted gene families using Ontologizer.

## Acknowledgement

We would like to thank Ren-jie Wang for sharing the snake photo, and three anonymous reviewers for their valuable comments. This project was supported by the Thousand Young Talents Program funding, and a startup funding of Life Science Institute of Zhejiang University to Q. Z., and National Major Scientific and Technological Special Project for ‘Significant New Drugs Development’ during the Twelfth Five-year Plan Period (No. 2009ZX09102-217, 2009-2010) to W. Y. All the raw reads, genome assembly and annotation generated in this study are deposited under the BioProject # PRJNA314370 on NCBI.

## Author Contribution

Q.Z., W.Y., G.Y. conceived and supervised the project. B.L., P.Q., W.Z., Y.H. and X.S. provided and extracted the samples. Z.L. and B.W. performed the proteomic mass spectrometry-based experiment and data analysis. J.L., Z.W. and Y.Z. performed the genome assembly and annotation. J.L. and Q.L. designed and performed the identification of sex-linked scaffolds. P.Z. performed RNA-seq data analysis. Z.W., L.J. and Y.Z. performed the genome evolution analyses. Z.W., Y.Z., L.J. and B.Q. performed genes and gene family evolution analyses. Z.W. and L.J. performed the sex chromosome evolution analyses. Q.Z., Z.W., G.Z., G.Y. interpreted the results and wrote the manuscript. All of the authors read and approved the final manuscript.

